# Protamine protects against Vancomycin Induced Kidney Injury

**DOI:** 10.1101/2024.08.12.607677

**Authors:** Justin Shiau, Patti Engel, Mark Olsen, Gwendolyn Pais, Jack Chang, Marc H. Scheetz

## Abstract

**Introduction:** Vancomycin causes kidney injury by accumulating in the proximal tubule, likely mediated by megalin uptake. Protamine is a putative megalin inhibitor that shares binding sites with heparin and is approved for heparin overdose in patients.

**Methods:** We employed a well characterized Sprague Dawley rat model to assess kidney injury and function in animals that received vancomycin, protamine alone, or vancomycin plus protamine over 5 days. Urinary KIM-1 was used as the primary measure for kidney injury while iohexol clearance was calculated to assess kidney function. Animals had samples drawn pre-treatment to serve as their own controls. Additionally, since protamine is not a known nephrotoxin, the protamine group also served as a control. Cellular inhibition studies were performed to assess the ability of protamine to inhibit OAT1, OAT3, and OCT2.

**Results:** Rats that received vancomycin had significantly increased urinary KIM-1 on day 2 (24.9 ng/24h, 95% CI 1.87 to 48.0) compared to the protamine alone group. By day 4, animals that received protamine with vancomycin had urinary KIM-1 amounts that were elevated compared to protamine alone (KIM-1 29.0 ng/24h, 95% CI 5.0 to 53.0). No statistically significant differences were identified for iohexol clearance changes between drug groups or when comparing clearance change from baseline (P>0.05). No substantial inhibition of OAT1, OAT3, or OCT2 was observed with protamine. IC_50_ values for protamine were 1e-4 M for OAT1 and OAT3 and 4.3e-5 M for OCT2.

**Conclusion:** Protamine, when added to vancomycin therapy, delays vancomycin induced kidney injury as defined by urinary KIM-1 in the rat model by one to three days. Protamine putatively acts through blockade of megalin and does not appear to have significant inhibition on OAT1, OAT3, or OCT2. Since protamine is an approved FDA medication, it has clinical potential as a therapeutic to reduce vancomycin related kidney injury; however, greater utility may be found by pursuing compounds with fewer adverse event liabilities.

## INTRODUCTION

Vancomycin is one of the most widely used antibiotics within United States in-patient healthcare systems, accounting for 11.9% of all antibiotic use (1,258 out of 10,612) in a 2015 hospital survey (1). As a tricyclic glycopeptide, it is primarily indicated for severe Gram-positive bacterial infections including methicillin-resistant *Staphylococcus aureus* (2). For about 70 years now, vancomycin has remained a gold-standard antibiotic widely used and studied across people of all ages. Despite its efficacy in the treatment of severe infections, one common limitation of its use is the association with acute kidney injury (AKI) (3). A systematic review and meta-analysis found that vancomycin is associated with a higher risk of AKI up to 2.5 times greater compared to other antibiotics (4).

Vancomycin kidney injury is attributed to two primary mechanisms, intracellular accumulation and complex formation with uromodulin and blockade of the proximal tubule (5, 6). While the individual contributions of each mechanism to the extent of kidney injury is not yet clear, intracellular accumulation is a hallmark (3, 6–8). Cellular accumulation occurs through multiple pathways both basolatereally and apically (6). Basolaterally, vancomycin is actively transported from blood circulation into kidney proximal tubule cells through the organic cation transporter (OCT)-2, located on the basolateral membrane of the tubular cell. Next, it is secreted into the tubular lumen from the apical membrane of the proximal tubule by an efflux transporter, P-glycoprotein (9, 10). Vancomycin that is in the tubular lumen (either via secretion or free filtration) is then transported across the apical membrane by apical endocytosis through dehydropeptidase (DHP)-1 and megalin. Intracellular vancomycin is aggregated by lysosomes leading to cascade of oxidative stress, complement activation, inflammatory injury, mitochondrial dysfunction, and cellular apoptosis (2).

Preventing vancomycin induced kidney injury is possible via multiple pathways. First, a very clear exposure response relationship exists (11–16). Lower area under the curve exposures result in less toxicity. Second, it is theoretically possible to prevent vancomycin from accumulating intracellularly. We (17) and others (18, 19) have previously demonstrated that administering cilastatin, a putative megalin inhibitor, reduces kidney injury(6, 18). However, the effect from cilastatin is modest and multiple mechanisms of action such as organic anion transporter inhibitions(20, 21) complicate mechanistic translation. Megalin is a large glycoprotein expressed on the apical surface of the proximal tubule that mediates intracellular signal transduction and plays an important role in the reabsorption of glomerular-filtered substances (18, 22). Demonstrating premise, megalin knock out mouse models have strong efficacy for suppressing drug induced kidney injury (23, 24). Vancomycin binds to megalin receptors which causes reuptake into these proximal tubule epithelial cells. Overall, transporter inhibition during active tubular secretion can have varying effects on drug concentrations (25–27). Inhibiting transporters remain an area of study that have potential to improve nephrotoxic drug administration.

Based on chemical structure, we hypothesized that protamine might block megalin and decrease vancomycin induced kidney injury. While relatively little information is available on the megalin binding capacity of protamine, protamine shares binding sites with heparin (current target for FDA approved use) and megalin(28, 29). Supporting this possibility of megalin blockade for the purpose of decreasing cellular accumulation and toxicity, Nagai and colleagues found that protamine inhibited the uptake of gentamicin in cultured opossum kidney cells that expressed megalin and cubilin (30). Megalin antagonists that block tubular drug reabsorption would be a significant advance as no drugs are approved for this indication.

## MATERIALS AND METHODS

### Experimental Design and Animals

This animal model study was conducted at Midwestern University in Downers Grove, Illinois with all methods reviewed and approved by MWU Institutional Animal Care and Use Committee (IACUC #3080). Male Sprague Dawley rats (n=19; age 8-10 weeks; 268-304g; Envigo, Indianapolis IN, USA) underwent double jugular catheterization (31) (Fig 1a). Surgery was performed under Ketamine (100mg/kg) and Xyalzine (10mg/kg) anesthesia with Buprenorphine (0.02mg/kg) given post-operatively for pain management. Animals were allowed to recover 4 days before administration of any drug. Animals were housed in metabolic cages starting one day prior to Day 0 with access to food and water *ad libitum* for the duration of the study (Fig 1b).

**Figure 1.**
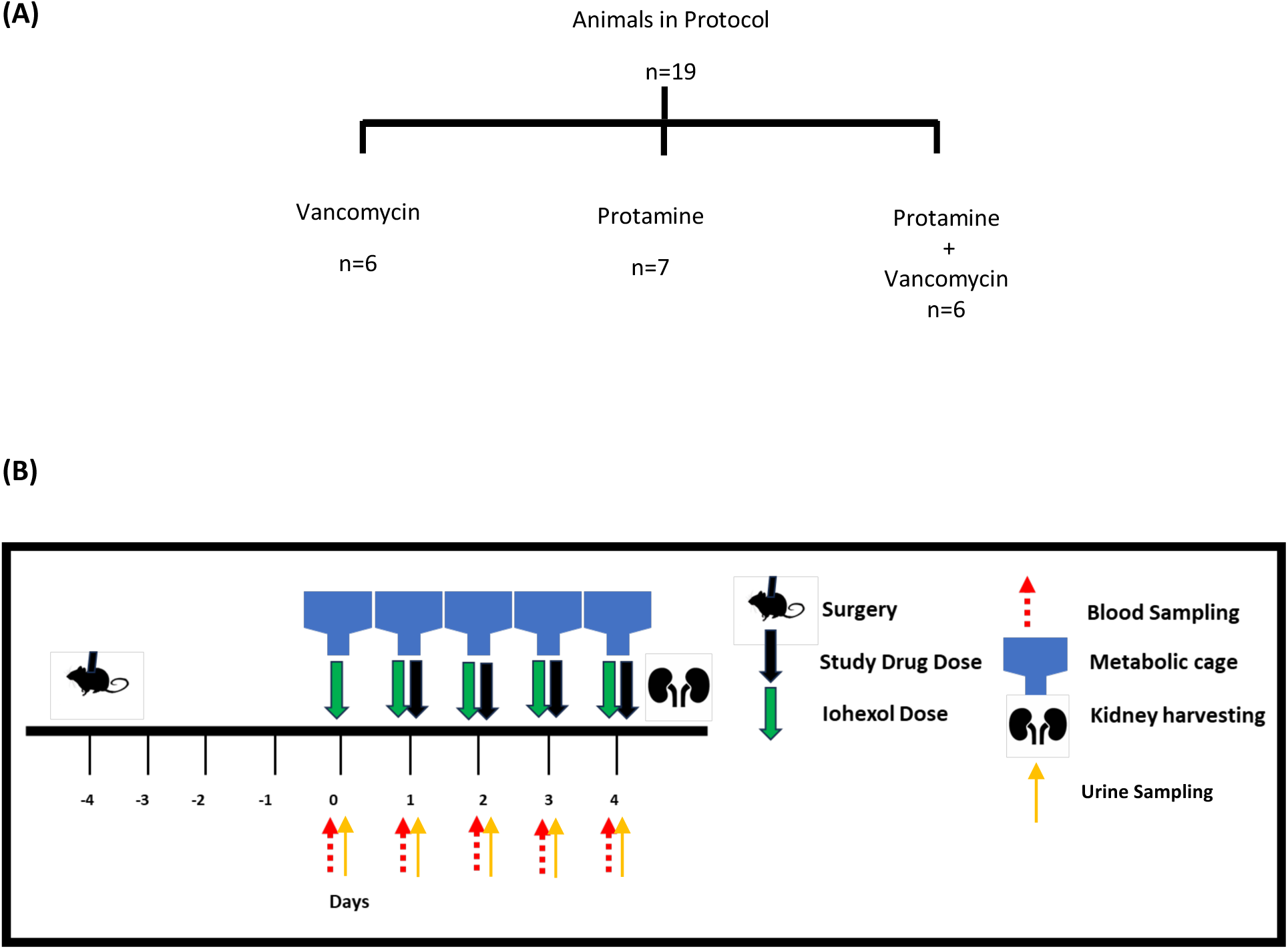
Experimental design **(A)** Animals divided into each protocol. **(B)** Scheme indicating time flow between surgery and when animals are placed into metabolic cages for dosing (PRT, VAN), blood and urine sampling and kidney harvest.

Animals were divided into 3 experimental treatment protocols. Using a dedicated jugular vein for dosing, the animals received clinical grade intravenous protamine and/or vancomycin in doses allometrically scaled to humans according to the following protocols: a) Protamine sulfate (1.5mg/kg) and Vancomycin (150mg/kg) (n=6); b) Protamine only (1.5mg/kg) with Saline SHAM (vancomycin) (n=7); or c) Vancomycin only (150mg/kg) with Saline SHAM (protamine) (n=6) (32–34). Saline SHAM was given to match volumes of protamine and vancomycin doses to maintain euvolemia. On Day 0, all animals received only Iohexol (51.8mg/day) intravenously to establish baseline GFR measurements. Drugs were administered as described above by treatment protocols to all groups on Days 1-4.

### Blood, Urine and Kidney Sampling

Blood samples were withdrawn from a designated jugular catheter. Samples were drawn at time 0 (prior to dosing) and at 30 and 60 minutes after Iohexol dosing for [protamine + SSHAM] and [vancomycin + SSHAM] groups. Sampling time points for [protamine + vancomycin] group were 0, 45 and 90 minutes to account for infusion times of both drugs and Iohexol in 6 animals. All blood samples (0.2ml, maximum 3 samples/animal) were taken daily with an equivalent volume of normal saline (NS) injected to maintain euvolemia. Blood samples (with EDTA added) were centrifuged at 16K x *g* 10 min. and resulting plasma was aliquoted (120*µ*l) and stored at −80C for batch analysis later.

Urine samples were collected daily, and volumes measured starting from Day 0 until Day 4. Samples were centrifuged at 500g x 10 minutes and the resulting supernatant aliquoted (1.5ml) and stored at - 80C for renal biomarker analysis later.

Animals were euthanized (Ketamine/Xylazine) on Day 4 after dosing/sampling for that day with terminal blood and urine collected. Kidneys were harvested, rinsed in cold NS; the right kidney was flash frozen in liquid nitrogen and stored at −80C. The left kidney was rinsed in NS then stored in 10% Buffered Formalin Phosphate and stored at room temperature.

### Chemicals and Reagents

Agents used for treatments were clinical grade Protamine Sulfate Injection (Fresnius Kabi, Lake Zurich, IL USA); clinical grade Vancomycin HCl (Hospira, Lake Forest, IL USA); Omnipaque (iohexol) injection (GE Healthcare, Marlborough, MA USA) and Normal Saline (Veterinary 0.9% sodium chloride injection USP; Abbott Laboratories, North Chicago, IL USA). For liquid chromatography – mass spectroscopy (LCMS) analysis LCMS grade acetonitrile, methanol and formic acid were used (VWR, Radnor, PA USA). Analytical grade iohexol (Sigma Aldrich, St. Louis, MO USA) and isotope Iohexol-d_5_ (Cayman Chemicals, Ann Arbor, MI USA) were also used. For the preparation of calibrators for standard curves, frozen aliquots of Na2EDTA treated (anticoagulant) non-medicated, non-immunized plasma pooled from male Sprague Dawley rats (BioIVT, Hicksville, NY USA) were used. For cellular experiments, chemical grade protamine sulfate USP (Product Number P3369, Sigma, St. Louis, MO) was used.

### Clearance as Calculated by Iohexol

In order to describe iohexol clearance a physiological PK model was created with non-linear mixed effects in Monolix (Version 2023R1; Antony, France: Lixoft SAS, 2023). A two-compartment model was used for the samples, with fixed estimates for peripheral compartment volume (V2) and intercompartmental transfer (Q) being derived from the base two-compartment model fitting. Random effects were estimated for clearance (Cl) and central volume (V1). The treatment group was evaluated as a modifier of clearance. To capture daily clearance changes, each experimental day (Monday through Friday) was considered as a separate occasion with time-varying clearance. Clearance was calculated as an occasion every 24 hours to capture the dynamic changes in renal function.

## EXPERIMENTAL METHODS

### Urinary Biomarkers

Urinary samples were analyzed for KIM-1 using Milliplex MAP Rat Kidney Toxicity Magnetic Bead Panel 1 (EMD Millipore Corporation, Charles, MO USA) according to manufacturer’s directions and previously described (32, 34, 35) Briefly, neat urine aliquots were mixed with monoclonal antibody coated magnetic microbeads (Bead Panel 1) and run on a 96 well plate which also included a standard curve. Biomarker data was acquired, and concentrations were analyzed using Belysa Immunoassay Curve-Fitting Software V1 (MilliporeSigma, Darmstadt, Germany). The amount of urinary KIM-1 excreted was obtained by multiplying the urinary concentration by 24-hour urinary volume.

### Preparation of Calibration curves in rat plasma

Stock solutions of Iohexol and Iohexol-d_5_ were prepared using purified water at concentrations of 1mg/ml and 100*µ*g/ml respectively. Calibrators for a standard curve were generated by diluting iohexol stock solution with water to produce concentrations ranging between 1.0 and 100*µ*g/ml.

For each standard curve calibrator, 36*µ*l of blank rat plasma was mixed with an iohexol dilution (4*µ*l each). Iohexol-d_5_ (4*µ*l) was then added to each at a final concentration of 10*µ*g/ml as the internal standard. Each standard curve calibrator was then mixed with 456*µ*l of 0.1% Formic Acid in methanol, vortexed and centrifuged at 16K x *g* for 10 min. From the resulting supernatant, 100*µ*l was transferred to LCMS vials and analyzed.

### Sample preparation

Plasma samples obtained at 30, 45 and 60 min were diluted 1:10 with blank rat plasma with 4µl Iohexol-d_5_ added to each. Iohexol-d_5_ was also added to remaining rat plasma samples (40µl, undiluted). All samples were then mixed with 0.1% Formic Acid in methanol, vortexed and centrifuged at 16K x *g* for 10 min. with resulting supernatant transferred to LCMS vials for analysis.

### LCMS Methods

An Agilent 1260 Infinity II series liquid chromatography system was paired with an Ultivo triple quadrupole mass spectrometer (Agilent Technologies, Santa Clara, CA) and used to analyze plasma samples. A Kinetex Polar C18 analytical column was used (2.6µm, 50 x 2.1mm, part number PRD-634714; Phenomenex) and column temperature was maintained at 25°C. Mobile phases were 0.1% formic acid in water (mobile phase A) and acetonitrile (mobile phase B). An isocratic method was used to separate analytes with solvent ratios of 94% mobile phase A – 6% mobile phase B (0 to 0.50 min), 5% mobile phase A – 95% mobile phase B (0.51 to 2.0 min) and 94% mobile phase A – 6% mobile phase B (2.01 to 3.5 min) at a mobile phase flow rate of 0.8 ml/min. Multiple reaction monitoring (MRM) was used for analyte detection monitoring transitions of *m/z* 821.9 to 803.9 for iohexol and *m/z* 826.9 to 808.7 for Iohexol-d_5_.

### Cell Studies

OAT1, OAT3, and OCT2 inhibition studies with protamine sulfate were performed by Eurofins Scientific (St. Charles, Missouri, USA) as previously described (36, 37). Protamine sulfate was dissolved in DMSO to 1.E-02 M and tested concurrently with p-aminohipurate 10 uM (substrate) for = OAT1, ranitidine 10 uM (substrate) for OAT3 and metformin 100uM (substrate) for OCT2. A time of 30 minutes was used for pre-incubation, followed by 20 minutes of incubation with the substrate. CHOK1 cells were used in OAT1 and 3 experiments, and HEK cells were used in OCT2 experiments. Percent of inhibition was calculated vs. the percent of control. The IC_50_ value (concentration causing a half-maximal inhibition of the control value) was determined by non-linear regression analysis of the concentration-response curve from a Hill equation. Experiments were compared to referent IC_50_ results on file for OAT1 and 3 using probenicid and for OCT2 using verapamil as exemplar inhibitors.

### Statistics

Urinary KIM-1 concentrations and iohexol clearances over time were compared with a repeated measures mixed model, fit via restricted maximum likelihood to have results comparable to the ANOVA results. An omnibus test, i.e. joint tests (multi degree of freedom) of the interaction and main effects which is similar to the F test in ANOVA, was used initially (STATA 17.0 BE, StataCorp LLC, College Station, Texas). Only models with significant interaction at p<0.05 were selected for post-hoc analysis (i.e., identification of the difference between treatment groups and days). Margins were calculated for a full factorial of the variables, i.e., main effects for each variable and interactions. Referent groups were pre-treatment baseline values and protamine alone as a treatment, except where noted. Secondarily, to remain agnostic to outcome variable relationship over time (and assess treatment groups over time), locally weighted scatterplot smoothing (LOWESS) trendlines with 95% confidence intervals were generated. LOWESS graphs were produced in R version 4.2.2 using ggplot2. No data were excluded as outliers. All tests were two-tailed, a p<0.05 was required for statistical significance. Graphs for cellular studies were generated using GraphPad version 9.3.1 and were fit with a log(inhibitor) vs. normalized response function after log transformation.

## RESULTS

### KIM-1

The omnibus test was significant for drug group, day, and the interaction of drug and day (P<0.05). Further evaluation of individual drug groups by day (with Bonferroni corrections) revealed that on day 1 prior to any drug received all groups had similar KIM-1 (Table 1). On day 2 (i.e. 24 hours after drug administration), animals that received vancomycin had significantly increased KIM-1 (24.9 ng/24h, 95% CI 1.87 to 48.0) compared to the protamine alone group as the base. Animals that received protamine did not experience this increase on day 2. On day 3, animals that received protamine had a measured KIM-1 that was numerically similar to animals that received vancomycin only. By day 4, animals that received protamine with their vancomycin had KIM-1 amounts that were elevated compared to protamine alone as a base comparison (KIM-1 29.0 ng/24h, 95% CI 5.0 to 53.0). When comparing each animal group to itself (i.e. change from baseline of day 1), animals that received protamine only did not have a KIM-1 increase over the 4 days (P>0.99) indicating that it was a stable control over the time course of the experiment (Table 2). Animals that received vancomycin alone had elevated KIM-1 on days 2 and 3 (P<0.05) compared to baseline day 1. Animals that received protamine with their vancomycin had KIM-1 elevated compared to baseline day 1 on day 4.

**Table 1.** Comparison of KIM-1 across drug treatment groups, by day.

| Drug | Day | Contrast KIM-1 (ng/24h) | Bonferroni P value | [95% conf. interval] |
| --- | --- | --- | --- | --- |
| Vancomycin vs. Protamine alone | 1 | 0.72 | >0.99 | [-22.3 to 23.8] |
| Vancomycin vs. Protamine alone | 2 | 24.92 | 0.025 | [1.9 to 48.0] |
| Vancomycin vs. Protamine alone | 3 | 20.88 | 0.106 | [-2.2 to 43.9] |
| Vancomycin vs. Protamine alone | 4 | 9.00 | >0.99 | [-14.0 to 32.0] |
| Vancomycin + Protamine vs. Protamine alone | 1 | 0.10 | >0.99 | [-22.9 to 23.1] |
| Vancomycin + Protamine vs. Protamine alone | 2 | -0.22 | >0.99 | [-23.3 to 22.8] |
| Vancomycin + Protamine vs. Protamine alone | 3 | 13.53 | 0.868 | [-9.5 to 36.6] |
| Vancomycin + Protamine vs. Protamine alone | 4 | 29.01 | 0.008 | [5.0 to 53.0] |

**Table 2.** Comparison of KIM-1 across over days within treatment groups with baseline as comparator.

| Drug | Day | Contrast KIM-1<br>(ng/24h) | Bonferroni P<br>value | [95% conf.<br>interval] |
| --- | --- | --- | --- | --- |
| Protamine + Vancomycin | 2 vs. 1 | 0.29 | >0.99 | [-19.3 to 19.8] |
| Protamine + Vancomycin | 3 vs. 1 | 13.78 | 0.457 | [-5.8 to 33.3] |
| Protamine + Vancomycin | 4 vs. 1 | 29.19 | 0.001 | [8.5 to 49.9] |
| Protamine alone | 2 vs. 1 | 0.61 | >0.99 | [-18.9 to 20.2] |
| Protamine alone | 3 vs. 1 | 0.35 | >0.99 | [-19.2 to 19.9] |
| Protamine alone | 4 vs. 1 | 0.28 | >0.99 | [-19.3 to 19.8] |
| Vancomycin | 2 vs. 1 | 24.81 | 0.004 | [5.2 to 44.4] |
| Vancomycin | 3 vs. 1 | 20.50 | 0.033 | [0.9 to 40.1] |
| Vancomycin | 4 vs. 1 | 8.55 | >0.99 | [-11 to 28.1] |

### Iohexol Clearance

The omnibus test was only significant for drug (P=0.047). No statistically observed differences were identified in the Bonferroni corrected results for comparing between drug groups or when comparing clearance change from baseline (P>0.05).

### Cellular studies

Cellular study results are shown in Figure 3. No substantial inhibition of OAT1, OAT3, or OCT2 was observed. IC_50_ values for protamine were 1e-4 M for OAT1 and OAT3 and 4.3e-5 for OCT2. Compared to prototype drugs causing inhibition (i.e. probenecid for OAT1/3 and verapamil for OCT2), IC_50_ values were 62.5 and 58.8 times higher for protamine versus probenecid (i.e. IC50s of 1.6E-06 M, 1.7E-06 M) and 5.2 times higher for protamine versus verapamil (IC50 of 8.3E-06 M).

## DISCUSSION

Our results demonstrate that the addition of protamine to vancomycin provides a protective effect against vancomycin AKI and that the effect is likely not driven by OAT1, OAT3, or OCT2 inhibition. These results are important as protamine is currently an FDA approved medication and has the potential to decrease vancomycin AKI, especially when given immediately prior to vancomycin for short vancomycin courses. These results may warrant a clinical study; however, protamine is not the most well tolerated medication and has the liabilities of severe hypotension and anaphylactoid-like reactions (38). Additional work is needed to confirm the putative mechanism of activity, i.e. megalin blockade. Drug development for medications with fewer adverse effects will be important if mechanism can be clarified.

Originally derived from salmon fish sperm (salmine), protamine is a highly positively charged protein consisting of 32 amino acids. Currently, it is widely used as an effective agent to neutralize heparin using electrostatic bonds made between the cationic arginine groups of protamine, and the anionic heparin in a 1:1 ratio. (6) Specifically, protamine shares binding sites with heparin (current target for FDA approved use) and megalin.(28, 29) Our preliminary data demonstrates that protamine does not have significant inhibition on OAT1, OAT3, or OCT2 except at extremely high concentrations (Fig 3) and as compared to known inhibitors such as probenecid and verapamil. Protamine enters kidney epithelial cells and localizes in the cytoplasm via receptor mediated endocytosis, putatively through megalin, and has demonstrated prevention of gentamicin accumulation.(39) Gentamicin is known to share a megalin cell entry mechanism with vancomycin.(18, 22, 40) Further, rats and humans share identical megalin extracellular motifs/ligand binding sites, making the rat ideal for pre-clinical study.(41, 42) Our pilot studies with protamine demonstrate a full day of injury prevention and a delay of 72 hours before mean injury for vancomycin+protamine reaches the vancomycin peak seen after 1 day of dosing, though full dose ranging experiments are required to define the dynamic toxicity threshold. Since ∼60% of clinical vancomycin use is ≤2 days,(43, 44) delaying toxicity will lower DIKI. In humans, vancomycin DIKI (creatinine defined)(45) occurs between an interquartile range of 6-13 days,(46) and creatinine is known to lag behind GFR.(47)

Reabsorption of solutes from glomerular filtrate in the kidneys occurs primarily in the proximal renal tubule.(48) Megalin is an endocytic transmembrane protein of the low-density lipoprotein receptor-related protein 2 family (LRP2).(22) It is located in the apical membrane of proximal tubule cells and reabsorbs low molecular weight proteins (LMW) through endocytosis.(22) Beenken et al. used high-resolution cryo-electron microscopy to study LRP2 isolated from mouse kidney.(49) Endosomes with receptor/ligand complex release ligands intracellularly through lysosome interaction, however megalin is recycled back to the apical membrane to be used again. They were able to demonstrate that ligand binding and release is pH dependent within extracellular (pH 7.5) and endosomal (pH 5.2) environments. Structurally, at the cell surface megalin adopts an open conformation exposing the high affinity ligand binding site at pH 7.5. At endosomal pH 5.2, megalin is transformed into a closed conformation with low binding affinity, thus releasing the ligand. The endosome with megalin moves back to the apical membrane where at extracellular pH 7.5 it once again resumes the open, high affinity conformation to bind ligand. Vancomycin is known to bind to megalin and its accumulation within proximal tubule cells is nephrotoxic causing oxidative damage and apoptosis.(3) Vancomycin, a basic compound, is protonated at urinary pH 7.5 and would bind to high affinity sites on Megalin, being released intracellularly in the proximal tubule cell as acidic pH induces the low affinity conformation.

The exact mechanism of vancomycin AKI is not entirely known, but current studies state that damage is directly caused by intracellular accumulation of vancomycin^(7)^. Vancomycin utilizes drug transporters located on the basolateral membrane of the proximal renal tubule including the organic anion transporter (OAT-1 and OAT-3), as well as the organic cation transporter (OCT-2). From the cell studies completed for our research (see cell study figure), it appears that with protamine, there is little to no inhibition of these transporters at all. With OAT-1 and OAT-3, inhibition does not even reach 50% at 1e-4 molar concentrations of protamine. With OCT-2, there does appear to be very minor inhibition reaching at least 50% at similar concentrations. This likely means that protamine acts as a weak, potentially non-pharmacologically useful substrate and/or inhibitor at incredibly high concentrations.

With the evidence that protamine has little to no interactions with the basolateral membrane transporters OAT-1, OAT-3, and OCT-2, the question still remains of how protamine is interacting with vancomycin to reduce AKI – or at the very least delay the onset of AKI when co-administered with vancomycin. It is known from current studies that vancomycin binds directly to the megalin receptor located within the proximal tubule. Hori et al performed a study to evaluate the binding of megalin to vancomycin and cisplatin, as well as aminoglycosides and colistin. They utilized quartz crystal microbalance (QCM) analysis to assess the direct binding of megalin with these drugs, and found that megalin is bound specifically by gentamicin, colistin, vancomycin, and cisplatin. Furthermore, they found that the binding of all above drugs listed is competed by cilastatin.

In humans, VAN DIKI (creatinine defined)(45) occurs between an interquartile range of 6-13 days,(46) and creatinine is known to lag behind GFR.(47). Serum creatinine is the most commonly utilized biomarker to classify AKI in patients,(45, 50–52) yet, creatinine is non-sensitive, non-specific, and slowly reactive to kidney function as compared to GFR.(53) GFR can drop 50% before changes in SCr are detected.(54–56) As such, acute injuries can take a median of 6 days to be detectable with creatinine alone (57). Therefore this study focused on the pre-clinical urinary biomarker, KIM-1, that has been shown as sensitive and specific for vancomycin kidney injury.(6, 58) Vancomycin induced kidney injury with urinary KIM-1 in the rat model is detectable within a single calendar day (6, 32) and the model links to human outcomes.(14) Protamine demonstrates a full day of injury prevention and a delay of 72 hours before mean injury for vancomycin + protamine reaches the vancomycin injury peak seen after 1 day of dosing (<u>Fig 2</u>). Since ∼60% of clinical VAN use is ≤2 days,(43, 44) delaying toxicity with an adjunct like protamine could lower vancomycin induced kidney injury.

**Figure 2.**
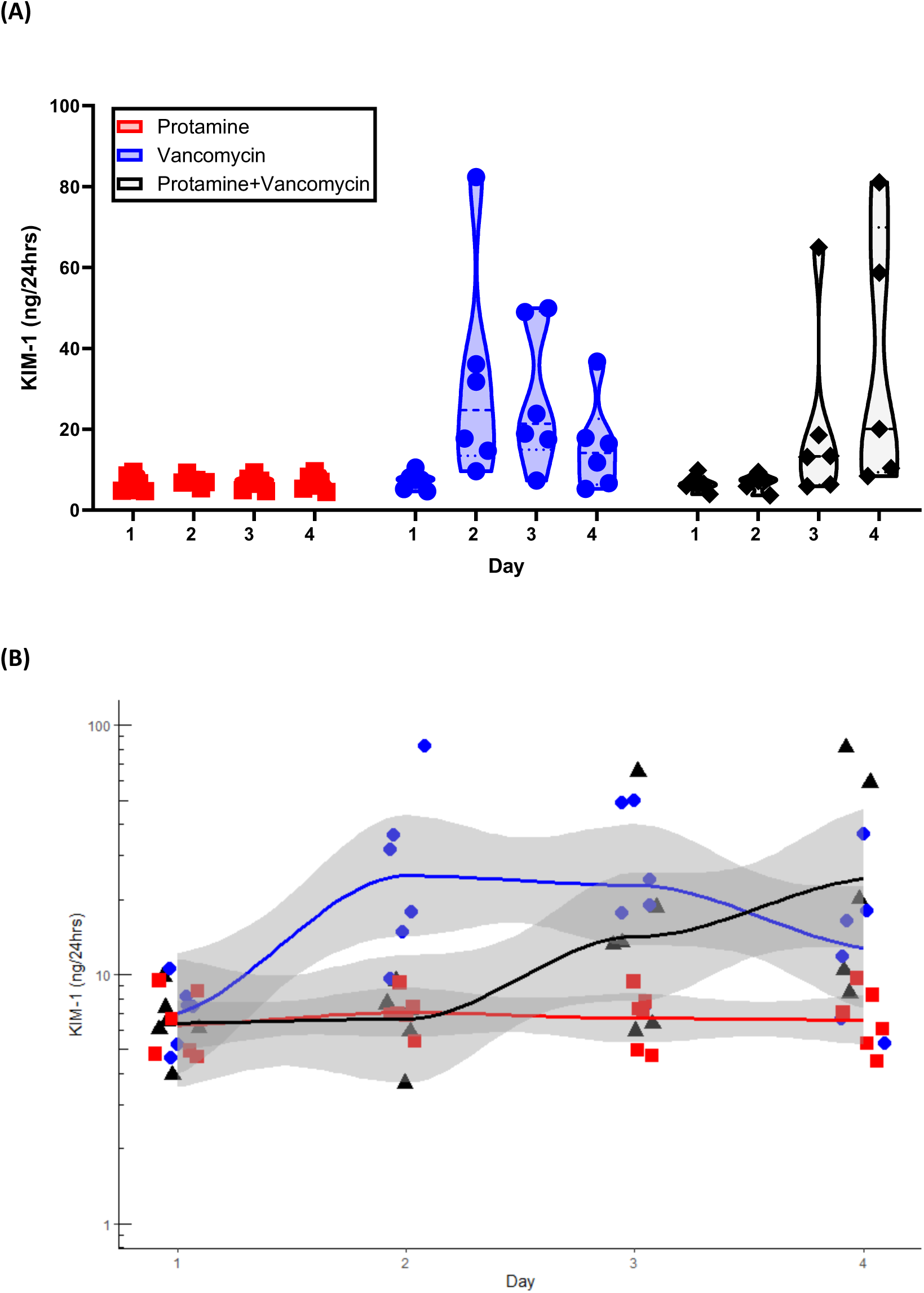
Urinary KIM-1 (ng/24) for each study group over study days, shown as a violin plot **(A)** and individual values with 95% CI **(B)**

**Figure 3.**
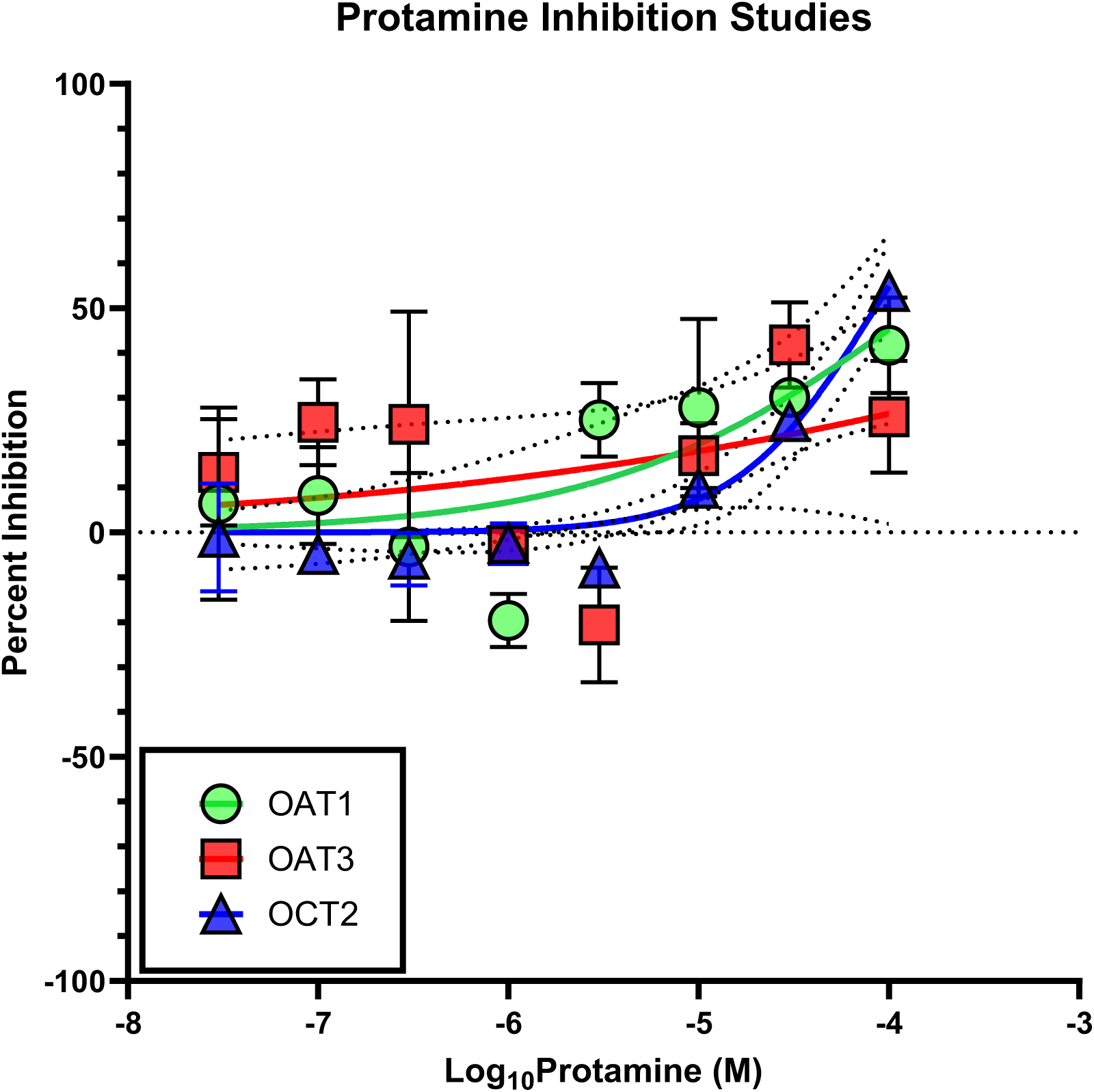
Cellular inhibition studies with protamine, shown in Molar (M) concentrations.

**Figure 4.**
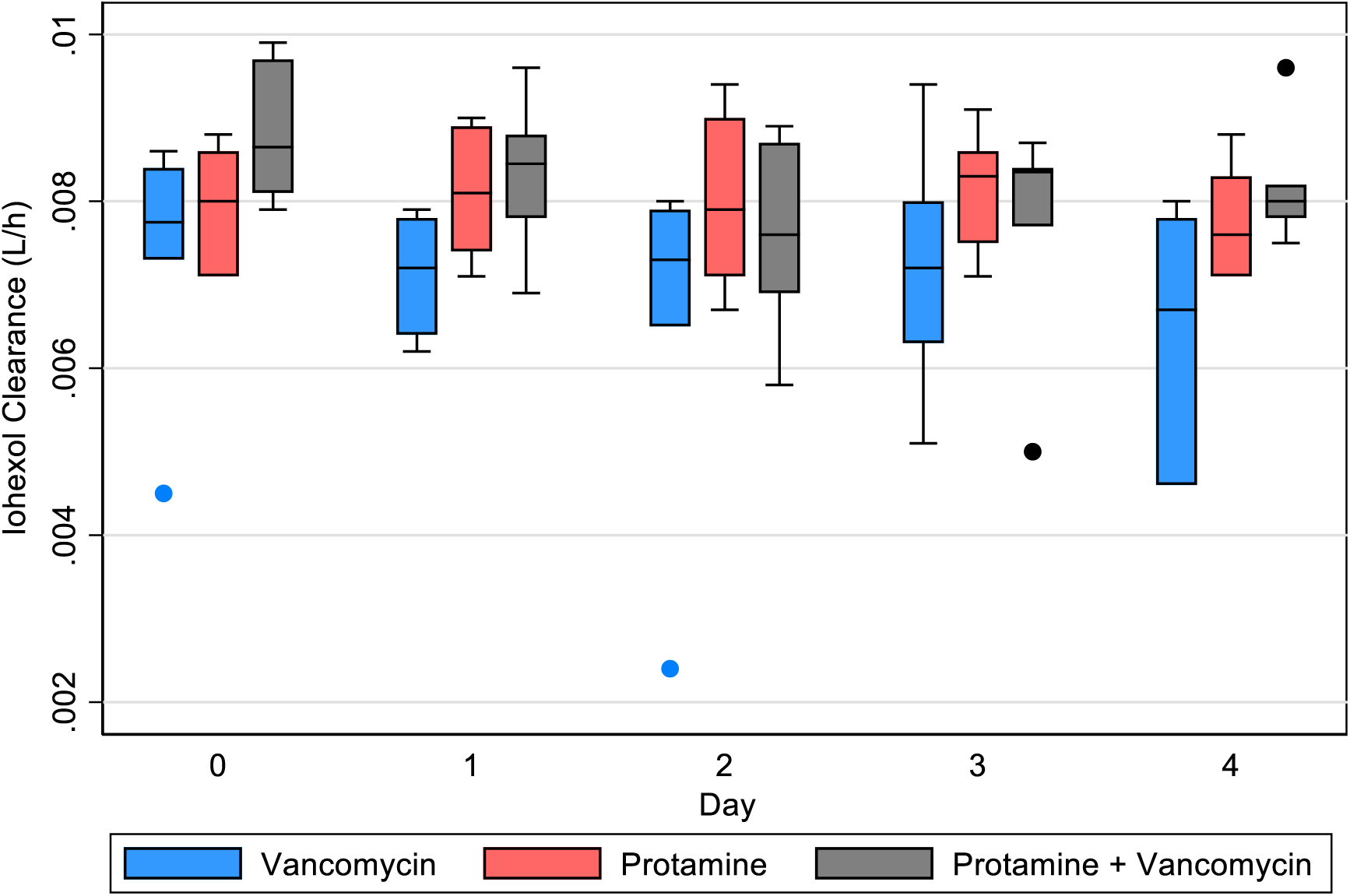
Calculated Iohexol clearance for each treatment group that received iohexol.

## CONCLUSION

Protamine, when added to vancomycin therapy, delays vancomycin induced kidney injury as defined by urinary KIM-1 in the rat model by one to three days. Protamine putatively acts through blockade of megalin and does not appear to have significant inhibition on OAT1, OAT3, or OCT2. Since protamine is an approved FDA medication, it has clinical potential as a therapeutic to reduce vancomycin related kidney injury; however, greater utility may be found by pursuing compounds with fewer adverse event liabilities.

